# Medial prefrontal cortical neurotransmitters reactive to relapse-promoting and relapse-suppressing cues in rats trained to self-administer cocaine or alcohol

**DOI:** 10.1101/2024.11.08.622733

**Authors:** Hermina Nedelescu, Cristina Miliano, Grant E. Wagner, Tony M. Kerr, Ann M. Gregus, Friedbert Weiss, Matthew W. Buczynski, Nobuyoshi Suto

## Abstract

Environmental cues conditioned to signal drug availability (S+) or omission (S-) activate specific neurons (neuronal ensembles/engram cells) within the medial prefrontal cortex (mPFC) to promote and suppress drug relapse in rats. However, the neurochemical source of such cue-specific activation remains unknown. In this study, we determined extracellular neurotransmitter fluctuations reactive to S+ vs. S- in the infralimbic (IL) and prelimbic (PL) cortices of male rats trained to lever-press for cocaine or alcohol self-administration. In cocaine- or alcohol-trained rats exposed to S+, no significant neurotransmitter fluctuations were observed in IL or PL. In cocaine-trained rats exposed to S-, glutamate, serotonin, taurine and adenosine were increased in PL but not in IL. In alcohol-trained rats exposed to S-, glutamate was increased, while dopamine and GABA were decreased, in IL but not in PL. Although S+ reactive neurotransmitters driving neuronal activation in mPFC remains to be elucidated, glutamate is likely the source of such activation by S- in rats trained to self-administer cocaine or alcohol. While drugs used for self-administration and cue-conditioning appear to dictate the type and anatomical specificity of S- evoked neurotransmission within mPFC, glutamate may serve as a common therapeutic target to mimic relapse-suppression by S- across cocaine and alcohol use disorders (CUD and AUD). In contrast, serotonin, taurine and adenosine may serve as the targets in CUD, while dopamine and GABA may serve as the targets in AUD.

## INTRODUCTION

Relapse-prevention is a major therapeutic goal in treating substance use disorders (SUDs). Environmental cues conditioned to signal drug availability (S+), along with stress and drugs of abuse themselves (drug priming), can trigger drug craving in recovering patients with SUDs and reinstate extinguished drug seeking in rats (Stewart, 2008). In contrast, environmental cues conditioned to signal drug omission (S-) can counter relapse-promotion by such drug availability cues, stress and drug priming in rats (Kearns et al., 2005; Mihindou et al., 2013; Suto et al., 2013; Suto et al., 2016; Laque et al., 2019; Madangopal et al., 2019; Hauser et al., 2023). Thus, rats trained to recognize both S+ and S- cues can serve as an animal model to determine brain mechanisms that either promote or suppress drug relapse, thereby aiding to identify neurobiological targets for anti-relapse medications.

Previous studies have found that S+ and S- each activate distinct functional units of neurons (neuronal ensembles/engram cells) in the infralimbic cortex (IL) – a ventral subregion of the medial prefrontal cortex (mPFC) – to promote and suppress relapse (Bossert et al., 2011; Suto et al., 2016; Warren et al., 2016; Laque et al., 2019; Warren et al., 2019). Such cue-specific neuronal activation could be mediated by cue-specific neurotransmission – a possibility that could directly translate into medications to either block or mimic the relapse-promoting vs. relapse-suppressing action of these cues. However, the neurochemical in the mPFC that are reactive to S+ and/or S- remain unknown.

The excitatory neurotransmitter glutamate (GLU) likely represents the primary neurochemical source of cortical activation. Neurons in IL are positioned to receive GLU from diverse cortical (e.g., prelimbic and orbitofrontal cortices) and subcortical (e.g., basolateral amygdala, hippocampus, thalamus and hypothalamus) brain regions (Hoover and Vertes, 2007) as well as from local pyramidal cells – the predominant neurons in the mPFC. Neurons in IL are also positioned to receive the inhibitory neurotransmitter γ-aminobutyric acid (GABA) from local GABAergic interneurons as well as dopamine (DA) from the midbrain (e.g., ventral tegmental area and substantia nigra compacta), serotonin (5-HT) from the raphe nucleus, norepinephrine (NE) from the locus coeruleus and acetylcholine (ACh) from the brainstem (e.g., pedunculopontine tegmental nucleus and laterodorsal tegmental nucleus). Additional small-molecule neurotransmitters are also present in the mPFC including adenosine (ADO), aspartate (ASP), D-serine (SER), glycine (GLY), histamine (HIS) and taurine (TAU). Since any of these neurotransmitters can regulate cortical neuron activity, we hypothesized that the relapse-promoting S+ and relapse-suppressing S- each engage each distinct combination of neurotransmitters in IL, ultimately contributing to their opposing actions on drug relapse.

Here, we used *in vivo* microdialysis to collect extracellular neurotransmitters from the IL and adjacent prelimbic cortex (PL) – the dorsal subregion of mPFC – similarly implicated in the regulation of drug relapse and cue-provoked drug seeking (Jonkman et al., 2009; Rocha and Kalivas, 2010; Pelloux et al., 2013; Stefanik et al., 2013). We collected dialysate samples from different groups of rats trained for operant cocaine or alcohol self-administration, representing two major classes of abused drugs as well as recognize two distinct odor stimuli (orange and almond scents) to discriminate between S+ and S- based on previously developed procedures (Suto et al., 2013; Suto et al., 2016; Laque et al., 2019). Since S+ and S- cues could affect any prefrontal cortical neurotransmitters, we implemented a high-throughput analytical method using high performance liquid chromatography-tandem mass spectrometry (HPLC-MS/MS) (Song et al., 2012; Zestos and Kennedy, 2017) to determine multiple small-molecule neurotransmitters in each dialysate sample. Specifically, we qualified and quantified the following 12 neurotransmitters in each sample: ADO, ASP, DA, GABA, GLY, GLU, GLN, HIS, NE, SER, 5- HT and TAU.

## MATERIALS AND METHODS

All animal/behavioral and histological procedures were conducted at The Scripps Research Institute (La Jolla, CA) in accordance with the National Institutes of Health (USA) Guidelines for the Care and Use of Laboratory Animals and approved by the local Institutional Animal Care and Use Committees. HPLC-MS/MS procedures were conducted at Virginia Tech (Blacksburg, VA). Data analyses were conducted at Scripps Research, Virginia Tech and Mayo Clinic (Rochester, MN).

### Subjects

Male Long Evans rats were used. All rats were purchased from Charles River, Inc. (Wilmington, MA). Rats weighing 200-250g for the “cocaine S+ vs. S-” experiment (see below) and 150-200g for the “alcohol S+ vs. S-” experiment (see below) at the start of experiments were housed in a temperature and humidity-controlled room, maintained on a 12 hour/12 hour reverse light/dark cycle. Water and food were available *ad libitum*. Rats assigned to the “alcohol S+ vs. S-” experiment were first acclimated to alcohol by receiving access to a drinking bottle containing 10% alcohol in home-cages for 3 weeks.

### Chemicals and Reagents

Cocaine was obtained from the Drug Depository Program from the National Institute on Drug Abuse (NIDA), National Institute of Health (NIH), USA. Alcohol was purchased from Sigma Aldrich (St. Louis, MO, USA). Chemicals for HPLC-MS/MS were purchased from Sigma (neurotransmitters, benzoyl chloride, benzoyl chloride-^13^C, sodium carbonate, sodium bicarbonate, ammonium formate, formic acid) and VWR (water, acetonitrile, methanol).

### Surgery

All rats were bilaterally implanted with microdialysis guide cannulae (Eicom, Kyoto, Japan). The guide cannula in one of the hemispheres was aimed at IL and the one in the other hemisphere was aimed at PL. Assignments of the left vs. right hemisphere for IL and PL placements were counterbalanced between rats. The coordinates for IL to place the tip of guide cannulae for microdialysis probes with 1.0 mm active membrane (see below) were +3.2 mm (posterior), ±0.5 mm (medial/lateral) and -4.4 mm (ventral) from the bregma. The coordinates for PL to place the tip of guide cannulae for microdialysis probes with 1.0 mm active membrane (see below) were +3.2 mm (posterior), ±0.5 mm (medial/lateral) and -2.8 mm (ventral) from the bregma. Rats assigned to the “cocaine S+ vs. S-” experiment were also implanted with intravenous catheters made of Micro-Renathane (Braintree Science, Braintree, MA, USA) as described previously (Laque et al., 2019). Rats recovered for 7 days.

### Behavioral Procedures

For all experiments, rats were trained and tested during the dark (active) phases in dedicated operant conditioning chambers (Med Associates, Saint Albans, VT, USA) based on previously developed procedures (Suto et al., 2013; Suto et al., 2016; Laque et al., 2019). Each chamber was equipped with two retractable levers (‘active’ and ‘inactive’ levers), a light-cue, a syringe pump, and a liquid swivel system for intravenous cocaine self-administration or *in vivo* microdialysis sample collection. The chambers used for the “alcohol S+ vs S-” experiment were also equipped with a drinking well for oral alcohol self-administration.

Both “cocaine S+ vs. S-” and “alcohol S+ vs S-” experiments consisted of five experimental phases:

i. Self-administration training: *The aim was to establish operant responding for cocaine or alcohol.* Rats were placed in operant conditioning chambers and habituated for 30 min. Insertion of active and inactive levers then began a once daily training session during which active lever-presses resulted in cocaine (1.0 mg/kg/infusion, intravenous) or alcohol (10% dissolved in water, 0.2 ml, oral) deliveries under a fixed ratio 1 (FR1) schedule of reinforcement. Each cocaine or alcohol delivery was paired with 20s or 2s light-cue illumination during which active lever-presses were recorded but had no scheduled consequences. Each training session for cocaine and alcohol lasted 240 min and 60 min, respectively. Rats underwent a minimum of 14 training sessions. Through this training, active lever and light-cue were conditioned to predict cocaine or alcohol 100%.
ii. Discrimination training: *The aim was to establish discriminative stimuli signaling cocaine or alcohol availability (S+) and omission (S-).* Rats were trained to recognize two olfactory cues (almond and orange odors) as S+ and S-. The odor assignment was counterbalanced between subjects. Each rat was subjected to alternating (twice daily) sessions to self-administer either cocaine or alcohol (preceded and accompanied by the S+ odor) or no cocaine or no alcohol (preceded and accompanied by the S- odor). Rats were placed and habituated in operant conditioning chambers for 30 min. The S+ or S- odor was then introduced 30 min prior to the insertion of two (active/inactive) levers and maintained throughout a 60-min session to lever-press for drug reward (cocaine/alcohol) or no reward (no cocaine/no alcohol). Each delivery of drug or no reward was paired with light-cue illumination (20s for cocaine/no cocaine and 2s for alcohol/no alcohol). Each rat underwent a minimum of 60 training sessions (30 each for S+ or S-). Through this training, each experimentally manipulated stimulus was conditioned to predict drug rewards (cocaine/alcohol) at the following probabilities: active lever and light-cue (50%), S+ odor (100%) and S- odor (0%).
iii. Discrimination tests for cocaine/alcohol seeking: *The aim was to test the behavioral effects of S+ and S-.* Each rat underwent three (once daily) cue-tests with the following conditions: [1] active lever and light-cue without S+ or S- (No S+/S- test), [2] active lever, light-cue and S+ (S+ test), [3] active lever, light-cue and S- (S- test). The cue-test order was counterbalanced between subjects. During all cue-tests, cocaine/alcohol was not available to isolate the effects of S+ or S- on operant response initiated and maintained by active lever and light-cue, as a measure of cue-provoked cocaine/alcohol seeking. For the No S+/S- test, rats were habituated in operant conditioning chambers for 60 min. Insertion of active/inactive levers began a 60 min cue-test to lever-press for light-cue illumination. For the S+ and S- tests, rats were habituated in operant chambers for 30 min. Next, the S+ or S- odor was introduced 30 min prior to inserting active/inactive levers and maintained throughout a 60-min session to lever-press for light-cue illumination.
iv. Discrimination re-training: *The aim was to re-establish S+ and S-.* Training procedures mimicked Phase II. Each rat underwent a minimum of 12 training sessions (6 each for S+ or S-).
v. Discrimination test for microdialysis: *The aim was to determine neurotransmitters reactive to S+ or S- in IL and PL.* On the day before this test, each rat received two microdialysis probes, one inserted into IL and the other into PL via surgically placed guide cannulae (see Surgery). We used the CX-I probes from Eicom Inc. (Kyoto, Japan) with 1.0 mm active membrane made of artificial cellulose with 50,000 Da molecular weight cut-off – suitable for broadly collecting small-molecular neurotransmitters. Rats were then placed and left in the operant conditioning chambers overnight equilibration; the probes were constantly perfused with artificial cerebrospinal fluid (aCSF) at 0.8 µl/min overnight and throughout the microdialysis procedures. On the following 2 days, rats underwent Discrimination test for microdialysis; each rat underwent two once daily cue-tests (S+ and S- tests). The order of S+ and S- tests were counterbalanced between subjects. Each cue-test was consisted of four experimental periods (30 min, each): [1] 30 min ‘baseline’ period without S+ or S- and active/inactive levers, [2] 30 min ‘cue-exposure’ period with S+ or S- but without active/inactive levers, [3] 30 min ‘operant responding’ period with S+ or S- and active/inactive levers and [4] 30 min ‘washout’ period without S+ or S- and active/inactive levers; microdialysis samples were collected at 10 min intervals. Rats were kept undisturbed for 30 min to collect the baseline microdialysis samples (baseline period). Next, rats were exposed to either S+ or S- odor for 30 min (cue-exposure period); active/inactive levers were not introduced during this period. Both active and inactive levers were then reintroduced, and the rats were given the opportunity to press these levers for 30 min while being exposed to either S+ or S- odor (operant responding period). In order to isolate the effects of environmental cues (S+/S- as well as active/inactive levers and light-cue illumination) from the pharmacological action of the drug (cocaine or alcohol), rats were tested under an extinction condition (i.e., cocaine and alcohol were not available during this test). The S+/S- odor and active/inactive levers were subsequently removed, and rats were kept undisturbed for 30 min to collect the “washout” samples (washout period).

After completing the first microdialysis experiment during Discrimination testing (S+ or S-), rats remained undisturbed in the same operant conditioning chamber overnight and underwent the second microdialysis experiment during Discrimination testing (S- or S+).. After completing both S+ and S- microdialysis testing experiments, rats were anesthetized with isoflurane and intracardially perfused with 100 ml of 1×phosphate buffered saline (PBS) followed by 200 ml of fixative solution (containing 4% paraformaldehyde and 14% saturated picric acid). Brains were collected and subsequently sectioned to histologically validate microdialysis probe placements.

### HPLC-MS/MS analysis

Quantification of neurotransmitters levels was performed using LC-MS/MS as previously described (Song et al., 2012; Buczynski et al., 2016). Each dialysate sample was used to quantify separately acetylcholine and all the other neurotransmitters. For the acetylcholine analysis, 2 µL of the sample was added to 5 µL ^4^d acetylcholine internal standard (in water) and acidified with 5 µL of formic acid (1%, water). For the neurotransmitter’s quantitation, 5 µL of the sample was added to 5 µL carbonate buffer (200 mM), derivatized with 2 µl of benzoyl chloride (2%, ACN), acidified with 2 µL of formic acid (1%, water), and supplemented 5µl internal standard (in ACN). Analysis was performed using a 1290 Infinity II LC System (Agilent Technologies) coupled with a 6495 triple quadrupole mass detector (Agilent Technologies) to measure, 5-HT (385→264), ASP (238→105), DA (466→105), GABA (208→105), GLU (252→105), GLN (251→105), HIS (216→105), NE (482→105), SER (210→105), TAU (230→105), adenosine (Ado, 372→136) and GLY (180→105). ^13^C benzoyl chloride-labeled counterparts were used as the internal standard. Although acetylcholine (ACh, 146→87) was included in the analytical method, it was not possible to detect it in the samples, therefore it was excluded from the analysis. Rats displaying unstable baselines were considered outliers and were removed from analysis.

### Statistical Analysis

Neurochemical data were analyzed separately for each neurotransmitter. Extracellular neurotransmitter concentrations were expressed as a percentage of baseline values: the mean concentration of the 6 samples collected during the 60-min ‘baseline’ period was defined as 100%. Each neurochemical was analyzed by using repeated measures ANOVA. For all cases, differences were considered significant when *P*<0.05 (two-tailed). When appropriate, ANOVA was followed by *post-hoc* tests. We used GraphPad Prism 8.4.3. (GraphPad, San Diego, CA, USA) or IBM SPSS Statistics 25 (IBM Corporation, Armok, NY, USA).

## RESULTS

### Infralimbic neurotransmitters reactive to S+ and S- in cocaine-trained rats

The infralimbic cortex plays a critical role in relapse-suppressing behavior, and our previous work shows that activated neuronal ensembles in this region during a cocaine-seeking paradigm. Thus, we first sought to evaluate neurochemical changes in the IFC that may underlie these molecular and behavioral events. Each neurotransmitter was evaluated by a separate two-way repeated measure ANOVA to determine the effect of time, cue (S+ vs S-), and the interaction between cue and time. We were surprised to discover that in cocaine-trained rats exposed to S+ or S-, there were no significant group or interaction changes in extracellular concentrations of any of the 12 neurotransmitters analyzed in the current study, including ADO, ASP, DA, GABA, GLY, GLU, GLN, HIS, NE, SER, 5-HT and TAU, in the dialysate samples collected from IL (**Table 1**). While HIS showed a modest time-dependent elevation throughout the experiment during S+ and S- (p < 0.05), these data do not support a role for this neurotransmitter in specific relapse-suppressing or relapse-promoting behaviors.

**Table 1.** Two-Way RM ANOVA Statistics for PL Dialysates from Cocaine IVSA Rats.

| NT | Brain Area | Factors | F-value | P-value |
| --- | --- | --- | --- | --- |
| ADO | PL | Cue x Time | F (11, 253) = 2.602 | <0.01 |
|  |  | Time | F (3.448, 79.30) = 1.345 | 0.264 |
|  |  | Cue | F (1, 23) = 4.011 | 0.057 |
| ASP | PL | Cue x Time | F (11, 209) = 0.7136 | 0.725 |
|  |  | Time | F (3.153, 59.91) = 0.3672 | 0.787 |
|  |  | Cue | F (1, 19) = 0.1632 | 0.691 |
| DA | PL | Cue x Time | F (11, 220) = 1.174 | 0.306 |
|  |  | Time | F (4.765, 95.30) = 1.236 | 0.299 |
|  |  | Cue | F (1, 20) = 0.1293 | 0.723 |
| GABA | PL | Cue x Time | F (11, 253) = 1.072 | 0.384 |
|  |  | Time | F (2.916, 67.07) = 0.5341 | 0.656 |
|  |  | Cue | F (1, 23) = 1.557 | 0.225 |
| GLN | PL | Cue x Time | F (11, 187) = 1.428 | 0.163 |
|  |  | Time | F (2.270, 38.59) = 2.121 | 0.128 |
|  |  | Cue | F (1, 17) = 0.5565 | 0.466 |
| GLU | PL | Cue x Time | F (11, 209) = 2.953 | <0.01 |
|  |  | Time | F (1.294, 24.59) = 4.038 | <0.05 |
|  |  | Cue | F (1, 19) = 3.241 | 0.088 |
| GLY | PL | Cue x Time | F (11, 231) = 0.6660 | 0.770 |
|  |  | Time | F (3.688, 77.46) = 0.7381 | 0.559 |
|  |  | Cue | F (1, 21) = 0.2650 | 0.612 |
| HYS | PL | Cue x Time | F (11, 253) = 0.4476 | 0.933 |
|  |  | Time | F (3.193, 73.44) = 1.787 | 0.154 |
|  |  | Cue | F (1, 23) = 0.2451 | 0.625 |
| NE | PL | Cue x Time | F (11, 231) = 0.7941 | 0.646 |
|  |  | Time | F (4.634, 97.32) = 0.7908 | 0.550 |
|  |  | Cue | F (1, 21) = 0.001247 | 0.972 |
| SER | PL | Cue x Time | F (11, 231) = 0.4566 | 0.928 |
|  |  | Time | F (2.847, 59.79) = 0.5437 | 0.645 |
|  |  | Cue | F (1, 21) = 0.02662 | 0.872 |
| TAU | PL | Cue x Time | F (11, 242) = 2.435 | <0.01 |
|  |  | Time | F (1.566, 34.46) = 2.457 | 0.112 |
|  |  | Cue | F (1, 22) = 4.019 | 0.057 |
| 5HT | PL | Cue x Time | F (11, 253) = 1.926 | <0.05 |
|  |  | Time | F (1.892, 43.51) = 1.426 | 0.251 |
|  |  | Cue | F (1, 23) = 2.152 | 0.156 |

### Prelimbic neurotransmitters reactive to S+ and S- in cocaine-trained rats

The prelimbic cortex plays a critical role in relapse-promoting behavior, and our previous work shows that activated neuronal ensembles in this region during a cocaine-seeking paradigm. Accordingly, each neurotransmitter from dialysate collected from the PL was evaluated by a separate two-way repeated measure ANOVA to determine the effect of time, cue (S+ vs S-), and the interaction between cue and time. We were again surprised to discover that in cocaine-trained rats exposed to S+, there were no significant group or interaction changes in extracellular concentrations of any of the 12 neurotransmitters analyzed in the current study, including ADO, ASP, DA, GABA, GLY, GLU, GLN, HIS, NE, SER, 5-HT and TAU, in the dialysate samples collected from PL (**Table 2**). In contrast, there was an interaction effect between S+ and S- indicating an elevation of GLU (**Fig 1A**, p<0.05) in the prelimbic cortex that was not present in the infralimbic cortex of these same rats (Fig 1B). Similar elevated responses were identified during the cue+lever phase in 5HT (p<0.05, **Fig 1C,D**), TAU (p<0.01, **Fig 1C,D**) and ADO (p<0.01, **Fig 1C,D**) during the cue+lever phase of the experiment in the prelimbic cortex. The remaining 8 neurotransmitters showed no significant changes in S+ or S- in the prelimbic cortex (**Table 1**) including ASP, DA, GABA, GLY, GLN, HIS, NE, and SER.

**Figure 1.**
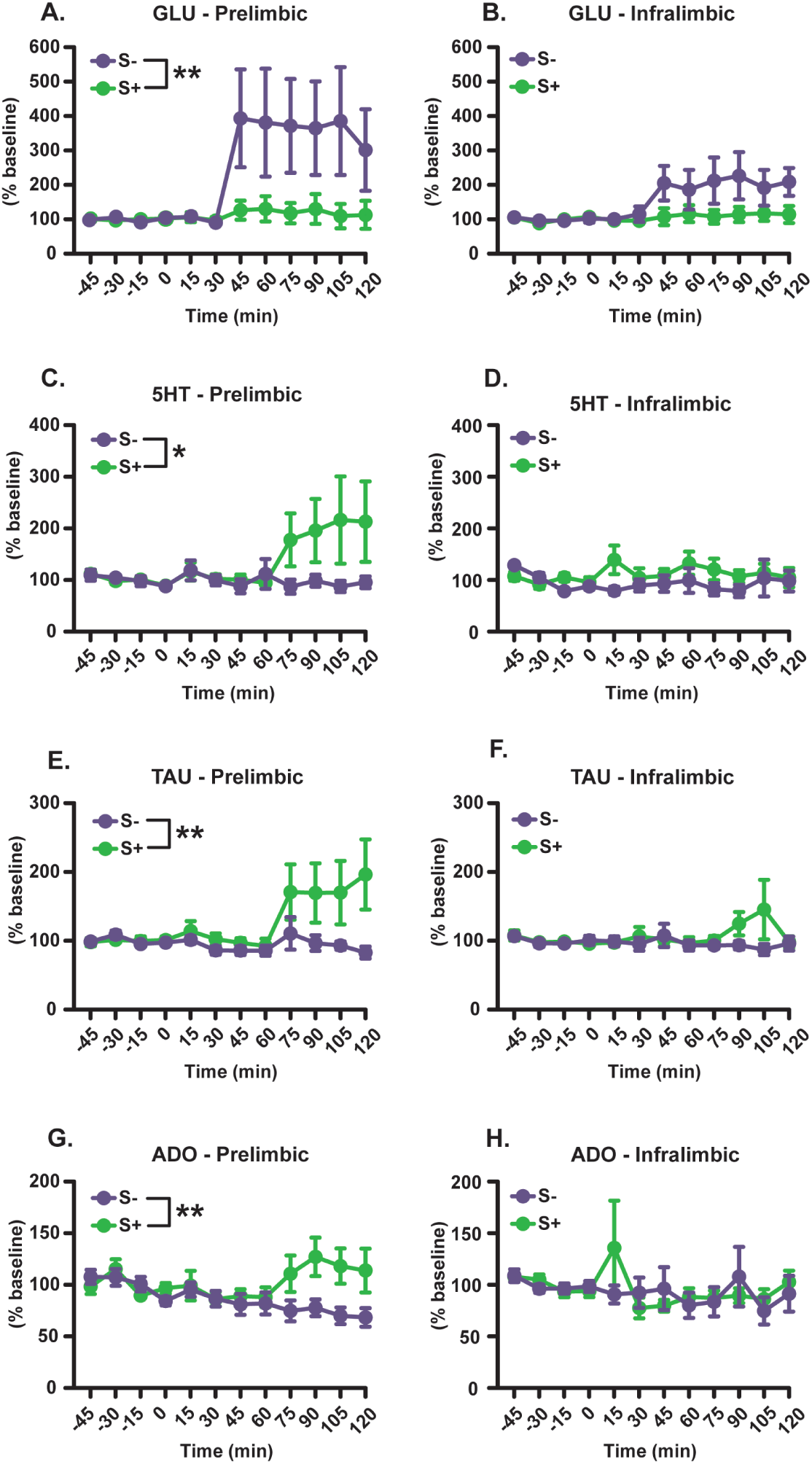
Neurochemical profiling of medial prefrontal cortex using *in vivo* microdialysis during cue discrimination in cocaine self-administering rats. Rats were trained to self-administer intravenous cocaine using an IVSA paradigm with both S+ and S- visual discrimination cues. Each rat was separately implanted with a microdialysis probe aimed at the prelimbic (left) and infralimbic (right) cortex and collected every 15 minutes during pre-session baseline (-45 to 0 min), introduction of the cue (0 to 30 min), and introduction of the cue-lever (30 to 120 min). Microdialysis samples were analyzed UPLC-MSMS for levels of 13 neurotransmitters and showed significant interaction effect (group x time) following two way RM-ANOVA between S+ and S- in the prelimbic cortex for **(A,B)** GLU, **(C,D)** 5-HT, **(E,F)** TAU, and **(G,H)** ADO. Data presented as mean ± SEM. Significance indicated by *p < 0.05, **p < 0.01.

**Table 2.** Two-Way RM ANOVA Statistics for IL Dialysates from Cocaine IVSA Rats.

| NT | Brain Area | Factors | F-value | P-value |
| --- | --- | --- | --- | --- |
| ADO | IL | Cue x Time | F (11, 264) = 0.7981 | 0.642 |
|  |  | Time | F (3.235, 77.64) = 1.111 | 0.352 |
|  |  | Cue | F (1, 24) = 0.06614 | 0.799 |
| ASP | IL | Cue x Time | F (11, 209) = 0.6429 | 0.791 |
|  |  | Time | F (3.068, 58.29) = 2.028 | 0.119 |
|  |  | Cue | F (1, 19) = 0.5194 | 0.480 |
| DA | IL | Cue x Time | F (11, 231) = 0.7956 | 0.644 |
|  |  | Time | F (6.546, 137.5) = 0.7625 | 0.611 |
|  |  | Cue | F (1, 21) = 1.293 | 0.268 |
| GABA | IL | Cue x Time | F (11, 253) = 0.7809 | 0.659 |
|  |  | Time | F (3.550, 81.64) = 0.7856 | 0.524 |
|  |  | Cue | F (1, 23) = 0.7825 | 0.386 |
| GLN | IL | Cue x Time | F (11, 165) = 0.5494 | 0.867 |
|  |  | Time | F (3.193, 47.89) = 0.4268 | 0.747 |
|  |  | Cue | F (1, 15) = 0.3476 | 0.564 |
| GLU | IL | Cue x Time | F (11, 176) = 1.690 | 0.079 |
|  |  | Time | F (2.166, 34.66) = 2.817 | 0.070 |
|  |  | Cue | F (1, 16) = 3.166 | 0.094 |
| GLY | IL | Cue x Time | F (11, 231) = 0.8428 | 0.597 |
|  |  | Time | F (2.029, 42.61) = 1.425 | 0.252 |
|  |  | Cue | F (1, 21) = 0.1022 | 0.752 |
| HYS | IL | Cue x Time | F (11, 242) = 1.077 | 0.380 |
|  |  | Time | F (5.687, 125.1) = 2.547 | <0.05 |
|  |  | Cue | F (1, 22) = 0.3578 | 0.556 |
| NE | IL | Cue x Time | F (11, 253) = 0.5394 | 0.876 |
|  |  | Time | F (3.691, 84.90) = 2.052 | 0.100 |
|  |  | Cue | F (1, 23) = 1.020 | 0.323 |
| SER | IL | Cue x Time | F (11, 242) = 0.6397 | 0.794 |
|  |  | Time | F (5.847, 128.6) = 0.5723 | 0.748 |
|  |  | Cue | F (1, 22) = 6.114e-005 | 0.994 |
| TAU | IL | Cue x Time | F (11, 242) = 1.297 | 0.227 |
|  |  | Time | F (2.054, 45.18) = 0.6567 | 0.527 |
|  |  | Cue | F (1, 22) = 1.335 | 0.260 |
| 5HT | IL | Cue x Time | F (11, 242) = 1.082 | 0.376 |
|  |  | Time | F (5.388, 118.5) = 0.7027 | 0.633 |
|  |  | Cue | F (1, 22) = 2.234 | 0.149 |

### Infralimbic neurotransmitters reactive to S+ and S- in alcohol-trained rats (*Fig 2*)

We performed a parallel study in rats trained to self-administer alcohol to evaluate the role of cue discrimination in a model of alcohol use disorder. Each neurotransmitter was evaluated by a separate two-way repeated measure ANOVA to determine the effect of time, cue (S+ vs S-), and the interaction between cue and time. In contrast with our cocaine experiment, alcohol-trained rats exposed to S+ or S- showed cue-dependent differences in extracellular GLU (p<0.05, Fig 2B) driven by elevations in the S- group. Additionally, alcohol-trained rats displayed cue-dependent differences in extracellular DA (p<0.05, Fig 2D) in the IL due to increased levels in the S- group. Finally, alcohol-trained rats showed cue-dependent differences in extracellular GABA (p<0.05, Fig 2F) that resulted from decreased levels in the S+ group. No significant changes in extracellular concentrations of any of the other 9 neurotransmitters analyzed including ADO, ASP, GLY, GLN, HIS, NE, SER, 5-HT and TAU in the dialysate samples collected from IL (**Table 3**).

**Figure 2.**
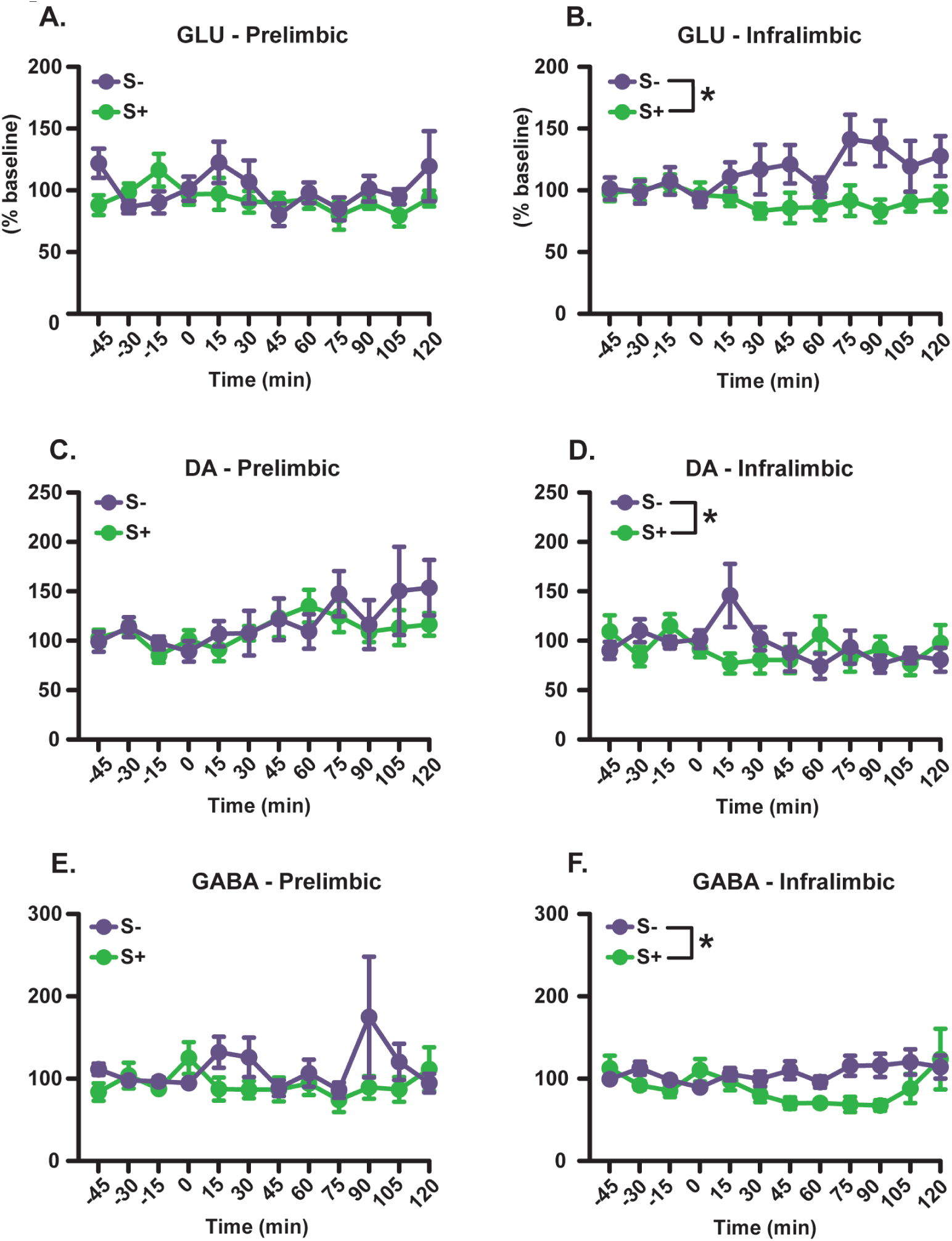
Neurochemical profiling of medial prefrontal cortex using *in vivo* microdialysis during cue discrimination in ethanol self-administering rats. Rats were trained to self-administer alcohol using S+ and S- visual discrimination cues. Each rat was separately implanted with a microdialysis probe aimed at the prelimbic (left) and infralimbic (right) cortex and collected every 15 minutes during pre-session baseline (-45 to 0 min), introduction of the cue (0 to 30 min), and introduction of the cue-lever (30 to 120 min). Microdialysis samples were analyzed UPLC-MSMS for levels of 13 neurotransmitters and showed significant group effect following two way RM-ANOVA between S+ and S- in the infralimbic cortex for **(A,B)** GLU, **(C,D)** DA, and **(E,F)** GABA. Data presented as mean ± SEM. Significance indicated by *p < 0.05.

**Table 3.**
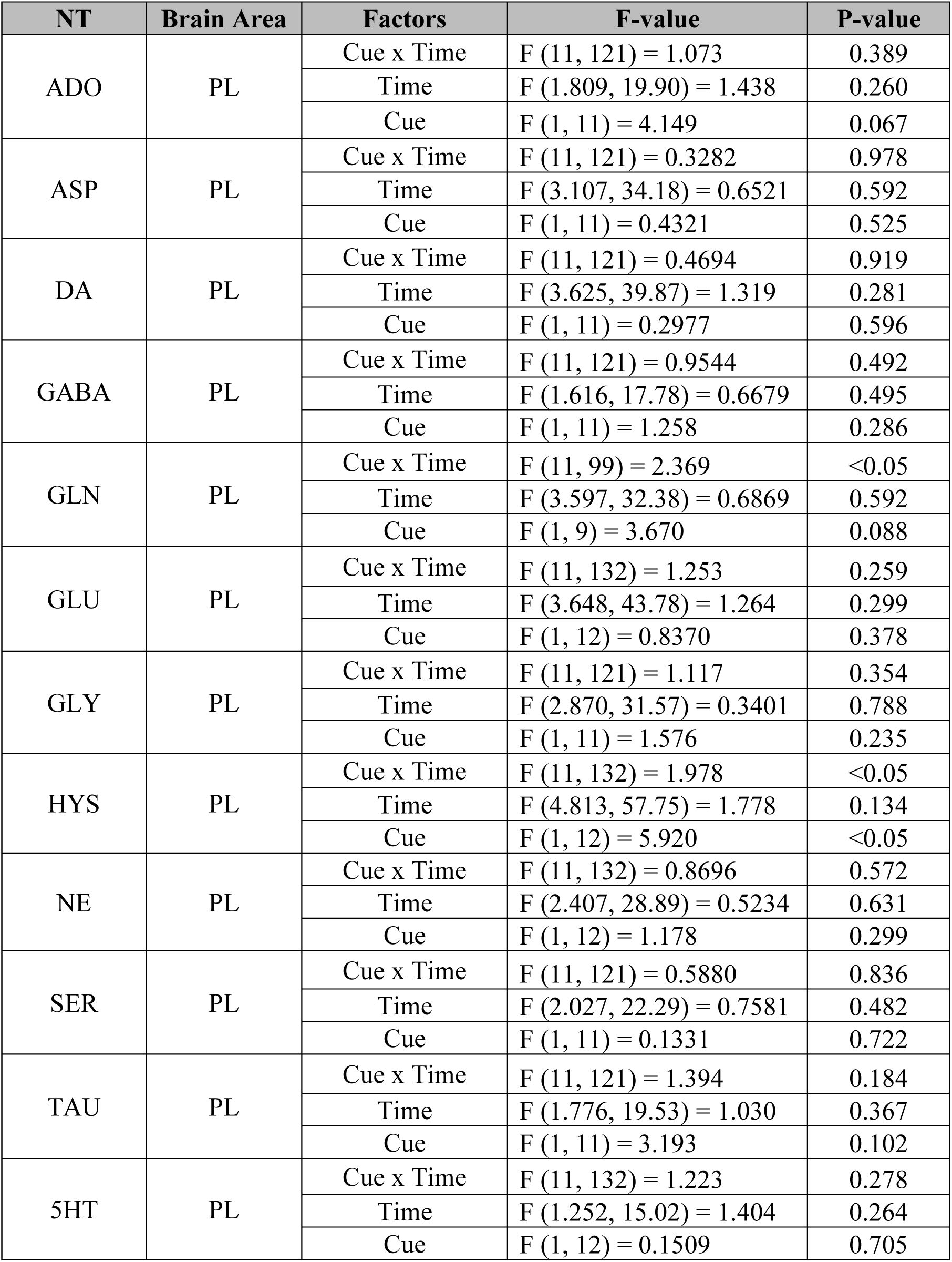
Two-Way RM ANOVA Statistics for PL Dialysates from Alcohol SA Rats.

### Prelimbic neurotransmitters reactive to S+ and S- in alcohol-trained rats

Finally, we performed an additional analysis in rats trained to self-administer alcohol to evaluate the role of cue discrimination on neurotransmitter levels in the prelimbic cortex. Accordingly, each neurotransmitter from dialysate collected from the PL was evaluated by a separate two-way repeated measure ANOVA to determine the effect of time, cue (S+ vs S-), and the interaction between cue and time. In contrast with our cocaine experiment, alcohol-trained rats exposed to S+ or S- exhibited no significant group or interaction changes in extracellular concentrations of any of the 12 neurotransmitters analyzed in the current study, including ADO, ASP, DA, GABA, GLY, GLU, GLN, HIS, NE, SER, 5-HT and TAU, in the dialysate samples collected from PL (**Table 4**).

**Table 4.** Two-Way RM ANOVA Statistics for IL Dialysates from Alcohol SA.

| NT | Brain Area | Factors | F-value | P-value |
| --- | --- | --- | --- | --- |
| ADO | IL | Cue x Time | F (11, 165) = 0.7756 | 0.664 |
|  |  | Time | F (1.124, 16.86) = 0.959 | 0.352 |
|  |  | Cue | F (1, 15) = 0.7184 | 0.410 |
| ASP | IL | Cue x Time | F (11, 143) = 0.7023 | 0.735 |
|  |  | Time | F (2.386, 31.02) = 3.251 | <0.05 |
|  |  | Cue | F (1, 13) = 0.6048 | 0.451 |
| DA | IL | Cue x Time | F (11, 165) = 2.107 | <0.05 |
|  |  | Time | F (5.500, 82.49) = 0.9745 | 0.443 |
|  |  | Cue | F (1, 15) = 0.2997 | 0.592 |
| GABA | IL | Cue x Time | F (11, 176) = 1.884 | <0.05 |
|  |  | Time | F (4.087, 65.39) = 1.311 | 0.275 |
|  |  | Cue | F (1, 16) = 5.490 | <0.05 |
| GLN | IL | Cue x Time | F (11, 143) = 1.392 | 0.183 |
|  |  | Time | F (3.626, 47.14) = 0.5644 | 0.673 |
|  |  | Cue | F (1, 13) = 0.8013 | 0.387 |
| GLU | IL | Cue x Time | F (11, 154) = 1.623 | 0.097 |
|  |  | Time | F (5.478, 76.70) = 0.7179 | 0.624 |
|  |  | Cue | F (1, 14) = 6.571 | <0.05 |
| GLY | IL | Cue x Time | F (11, 165) = 0.8206 | 0.620 |
|  |  | Time | F (2.921, 43.81) = 1.060 | 0.375 |
|  |  | Cue | F (1, 15) = 0.04305 | 0.838 |
| HYS | IL | Cue x Time | F (11, 165) = 0.4156 | 0.948 |
|  |  | Time | F (2.102, 31.53) = 2.192 | 0.126 |
|  |  | Cue | F (1, 15) = 0.2008 | 0.661 |
| NE | IL | Cue x Time | F (11, 154) = 0.6506 | 0.783 |
|  |  | Time | F (3.467, 48.54) = 0.6895 | 0.583 |
|  |  | Cue | F (1, 14) = 0.003986 | 0.951 |
| SER | IL | Cue x Time | F (11, 165) = 0.9278 | 0.515 |
|  |  | Time | F (1.289, 19.34) = 0.6595 | 0.464 |
|  |  | Cue | F (1, 15) = 0.5116 | 0.485 |
| TAU | IL | Cue x Time | F (11, 176) = 0.2648 | 0.991 |
|  |  | Time | F (1.900, 30.41) = 1.997 | 0.155 |
|  |  | Cue | F (1, 16) = 1.046 | 0.322 |
| 5HT | IL | Cue x Time | F (11, 165) = 1.812 | 0.056 |
|  |  | Time | F (1.636, 24.54) = 1.643 | 0.216 |
|  |  | Cue | F (1, 15) = 0.05992 | 0.810 |

## DISCUSSION

In the current study, we determined the extracellular fluctuations in S+ vs S- reactive small-molecule neurotransmitters in the IL and PL of male rats trained to lever-press for either cocaine or alcohol self-administration. We used *in vivo* microdialysis to collect neurochemical samples in the IL or PL from rats exposed to either S+ or S-, and subsequently given an opportunity to lever-press under an extinction condition in the absence of the primary drug reward (no cocaine or alcohol) as an animal model of drug-seeking behavior. As previously reported, rats previously trained to self-administer either cocaine or alcohol and recognize two distinct odor cues (orange and almond scents) as discriminative stimuli signaling drug availability and omission actively engaged in lever-pressing when exposed to S+ (but not S-) under an extinction condition (Weiss et al., 2001; Martin-Fardon and Weiss, 2017). Previous studies have also established that such S- as drug omission cue can actively suppress the relapse-promoting actions of not only drug availability cues but also stress and drug itself (priming) (Kearns et al., 2005; Mihindou et al., 2013; Suto et al., 2013; Suto et al., 2016; Laque et al., 2019; Madangopal et al., 2019; Hauser et al., 2023). Since S+ and S- could modulate any prefrontal cortical neurotransmitters, we used a high-throughput analytical method (Song et al., 2012; Zestos and Kennedy, 2017) to qualify and quantify the following 12 neurotransmitters known to be present in the mPFC: ADO, ASP, DA, GABA, GLY, GLU, GLN, HIS, NE, SER, 5- HT and TAU.

In cocaine- or alcohol-trained rats exposed to S+, no significant extracellular fluctuations in the 12 neurotransmitters analyzed were observed in IL or PL (Figure 1 and 2) despite these rats actively engaging in lever-pressing behavior. Importantly, this includes no significant changes in the major excitatory neurotransmitter GLU and another excitatory neurotransmitter ASP. While the neuroanatomical specificity and functional significance of the IL and PL in such cue-provoked drug seeking remains controversial (Moorman et al., 2015; Gourley and Taylor, 2016; Suto et al., 2016; Moorman and Aston-Jones, 2023), the current results are surprising considering the extensive literature implicating medial prefrontal cortical neuronal activity in cue-provoked drug seeking (Willcocks and McNally, 2013; Moorman and Aston-Jones, 2015; Pfarr et al., 2015; Warren et al., 2019). Indeed, previous reports also implicate medial prefrontal cortical GLU (Park et al., 2002; Kalivas et al., 2003; Van den Oever et al., 2008; Schmidt and Pierce, 2010; Parsegian and See, 2014), DA (McGlinchey et al., 2016; James et al., 2018), GABA (Van den Oever et al., 2010; Chefer et al., 2011; Lubbers et al., 2014) and 5-HT (Anastasio et al., 2014; Swinford-Jackson et al., 2016) neurotransmission in the ‘reinstatement’ of cue-provoked drug seeking. However, these studies largely did not measure extracellular neurotransmitter fluctuations using microdialysis or other direct *in vivo* neurochemical monitoring methods, such as voltammetry and biosensors, and instead relied on pharmacological manipulations to establish the roll of different transmissions in cue-provoked drug seeking.

While the discrepancies between these prior studies and our current findings need to be fully addressed, a similar lack of cue-evoked GLU in the mPFC that accompanied cue-provoked cocaine seeking was observed 3 days after the last cocaine self-administration (Shin et al., 2016). The lack of cue-evoked GLU during ‘early’ drug abstinence is consistent with the current microdialysis results collected 2 days after the last cocaine or alcohol self-administration (Figure 2). In contrast, the same study identified a significant increase in cue-evoked GLU 30 days after the last cocaine self-administration accompanied by significantly enhanced cue-provoked drug seeking (Shin et al., 2016). Moreover, previous studies implicating medial prefrontal cortical glutamatergic transmission in the ‘reinstatement’ of cue-provoked drug seeking (Park et al., 2002; Kalivas et al., 2003; Van den Oever et al., 2008; Schmidt and Pierce, 2010) were also conducted during similarly ‘protracted’ abstinence following extinction training that typically lasted for several weeks. Thus, glutamatergic transmission in the mPFC appears to mediate the expression of cue-provoked drug seeking during protracted (several weeks), rather than early (several days), drug abstinence. Consistent with this hypothesis, pharmacological manipulation of – either increased or decreased – glutamatergic transmission in the PL or IL significantly altered cue-provoked cocaine seeking 30 days after – but not 3 days after – the last cocaine self-administration (Shin et al., 2018).

Nevertheless, previous studies have implicated cue-provoked medial prefrontal cortical activation (presumably mediated by local excitatory neurotransmission) in cue-provoked drug seeking during early drug abstinence (Moorman and Aston-Jones, 2015, 2023). Since S+ did not affect the excitatory neurotransmitters GLU and ASP in IL or PL (Figure 1) despite provoking active lever-pressing in the current study, the neurochemical source of such activation remains to be determined. We speculate here that the excitatory neurotransmitter acetylcholine (ACh) represents one of the possible sources of cue-provoked neuronal activation in mPFC during early drug abstinence. Indeed, ACh is present in mPFC (Lacroix et al., 2006; Huang et al., 2014; Fecik et al., 2024) and implicated in both cue-provoked drug seeking (Schmidt et al., 2009) and drug cue-mediated conditioned place preference (Pastor et al., 2021). Unfortunately, while we attempted to analyze ACh without the use of cholinesterase inhibitors in dialysates (Song et al., 2012; Zestos and Kennedy, 2017), our adaptation of this method proved to be insufficient perhaps due to technical differences in the instrumentation as well as sampling methods and location.

Despite the lack of active lever-pressing, cocaine-trained rats exposed to S-, multiple neurotransmitters, including GLU, 5-HT, TAU and ADO, were significantly increased in PL but not IL (Figure 2). It is worth noting that the S- evoked increase in GLU overflow in IL (Figure 2) approached significance (P = 0.050) in these cocaine-trained rats. In alcohol-trained rats exposed to S-, GLU and DA were increased while GABA was decreased in IL, but not PL (Figure 2). Taken together, these results indicate that the suppression of drug seeking by a discriminative stimulus conditioned to signal drug unavailability or omission (Kearns et al., 2005; Mihindou et al., 2013; Suto et al., 2016; Laque et al., 2019; Madangopal et al., 2019; Hauser et al., 2023) is mediated by active neurochemical mechanisms in the brain. In turn, this raises the possibility of using S- reactive neurotransmitters as a therapeutic target to develop medication that mimics the relapse-suppressing action of drug omission cues.

Previous studies have found that S- activates distinct neuronal units (neuronal ensembles/engram cells) in the IL to suppress drug and non-drug reward seeking (Suto et al., 2016; Warren et al., 2016; Laque et al., 2019; Warren et al., 2019). Moreover, previous studies also implicate PL neuronal activation in suppressing of drug seeking (Moorman and Aston-Jones, 2015, 2023). While drugs used for self-administration and cue-conditioning appear to dictate the type and anatomical specificity of S- evoked neurotransmission within the mPFC, our current results indicate that GLU is the likely the source of activation by S- in rats trained to self-administer cocaine or alcohol. Thus, glutamatergic transmission in mPFC may serve as a common therapeutic target to mimic relapse-suppression by S- across cocaine and alcohol use disorders (CUD and AUD) regardless of the anatomical IL and PL sources. Indeed, CUD and AUD are both associated with hypoactive prefrontal cortex – a physiological condition linked to diminished executive control and heightened relapse risk (Goldstein and Volkow, 2002; Vetreno and Crews, 2014). Thus, drugs which can selectively target to mimic S- evoked GLU overflow and enhance glutamatergic transmission in the mPFC may prove to be beneficial for treating cue-provoked drug seeking and craving in AUD and CUD.

Since S- increased prelimbic 5-HT, ADO and TAU overflow selectively in cocaine-trained (vs. alcohol-trained) rats, these three neurotransmitters may serve as the therapeutic targets in CUD. Indeed, 5-HT receptor agonists such as lorcaserin have been shown to be moderately effective at reducing cocaine seeking in rats (Anastasio et al., 2020) and cocaine-choice in humans (Pirtle et al., 2019; Grasing et al., 2022). Moreover, ADO receptor agonists have shown promising preclinical results against cocaine motivated behavior (Ballesteros-Yanez et al., 2017; Borroto-Escuela et al., 2018). While the therapeutic action of taurine in CUD has not been extensively investigated, the derivative of its endogenous analogue homotaurine acamprosate – an FDA approved medication for AUD (Heilig, 2014; Mason, 2022) – has demonstrated efficacy at reducing cue-provoked cocaine seeking in rats (Bowers et al., 2007), although a pilot clinical trial of acamprosate in patients with CUD produced negative results (Kampman et al., 2011). Nevertheless, the chemical structure of taurine is similar but distinct from homotaurine or acamprosate, and thus it may be beneficial to determine taurine’s therapeutic efficacy in CUD. In principle, pharmacologically mimicking S- evoked increase in 5- HT, ADO and TAU neurotransmission may similarly mimic the relapse-suppressing action of S- on cue-provoked cocaine seeking in rats and facilitate relapse-prevention in patients with CUD.

In contrast, since S- decreased infralimbic DA and GABA overflow only in alcohol-trained (vs. cocaine-trained) rats, mimicking these neurotransmissions could mimic the relapse-suppressing action of S- on cue-provoked alcohol seeking in rats and potentially help prevent relapse in patients with AUD. However, since reduced prefrontal cortical DA levels is linked to diminished cognitive function and increased alcohol relapse risk (Trantham-Davidson and Chandler, 2015), it is unclear if mimicking the S+ evoked decrease in infralimbic dopaminergic transmission can suppress cue-provoked alcohol seeking and craving. Nevertheless, dopaminergic antagonists such as tiapride, clozapine and ondansetron, have all demonstrated mild efficacy in reducing alcohol craving and consumption in clinical trials, although the therapeutic actions of these compounds were plagued with dopaminergic side-effects (Swift, 2010). Since the FDA approved anti-AUD medication acamprosate is known to act as a positive allosteric modulator of GABA-A receptors, it is also not clear that mimicking S- evoked decrease in infralimbic GABAergic transmission could suppress cue-provoked alcohol seeking and serve as a therapeutic treatment for AUD. However, the exact mode of therapeutic action of acamprosate in AUD is still not clear; some findings suggesting the anti-relapse action of acamprosate is mediated by inhibition of N-methyl-D-aspartate GLU receptors rather than positive allosteric modulation of GABA-A receptors (Heilig, 2014; Mason, 2022), while others suggest it is mediated by calcium (Spanagel et al., 2014). Taken together, further research is clearly needed to fully establish the therapeutic potential of mimicking S+ evoked changes in prefrontal cortical DA and GABA to facilitate relapse-prevention in AUD.

## ACKNOWLEDGEMENTS

This work was supported by Extramural and Intramural funding from National Institute on Drug Abuse, National Institute of Alcohol Abuse and Alcoholism and National Cancer Institute, National Institute of Health, USA: K01DA054449 (H.N.), R01DA037294 (N.S.), R01AA023183 (N.S.), U01DA055017 (N.S.), R21AA030184 (N.S.), R01CA284075 (M.W.B), R01AA027555 (F.W.), R01AA014351 (F.W.) and R01AR075241 (A.M.G.). H.N. was also supported by Ruth L. Kirschstein Institutional National Research Service Award from National Institute of Alcohol Abuse and Alcoholism, National Institute of Health, USA: T32AA007456 (PIs, Drs. Michael Taffe and Marisa Roberto).

## Notes

### Competing Interest Statement

The authors have declared no competing interest.

